# The influence of training-induced sarcomerogenesis on the history dependence of force

**DOI:** 10.1101/2020.03.31.018895

**Authors:** Jackey Chen, Parastoo Mashouri, Stephanie Fontyn, Mikella Valvano, Shakeap Elliott-Mohamed, Alex M. Noonan, Stephen H. M. Brown, Geoffrey A. Power

**Affiliations:** Department of Human Health and Nutritional Sciences, College of Biological Sciences, University of Guelph, Guelph, Ontario, Canada

**Keywords:** Fascicle, muscle, residual force enhancement, residual force depression, sarcomere, eccentric, concentric, uphill running, downhill running

## Abstract

The increase or decrease in isometric force following active muscle lengthening or shortening, relative to a reference isometric contraction at the same muscle length and level of activation, are referred to as residual force enhancement (rFE) and residual force depression (rFD), respectively. The purpose of these experiments was to gain further mechanistic insight into the trainability of rFE and rFD, on the basis of serial sarcomere number (SSN) alterations to length-dependent properties. Maximal rFE/rFD measures from the soleus and extensor digitorum longus (EDL) of rats were compared after 4 weeks of uphill/downhill running and a no running control. Serial sarcomere numbers adapted to the training: soleus serial sarcomere number was greater with downhill compared to uphill running, while EDL demonstrated a trend towards more serial sarcomeres for downhill compared to no running. In contrast, absolute and normalized rFE/rFD did not differ across training groups for either muscle. As such, it appears that training-induced SSN adaptations do not modify rFE/rFD at the whole-muscle level.

**Summary Statement:** The addition and subtraction of serial sarcomeres induced by downhill and uphill running, respectively, did not influence the magnitude of stretch-induced force enhancement and shortening-induced force depression.

## Introduction

The history dependence of force is an intrinsic property of skeletal muscle that has been investigated in individual sarcomeres (Joumaa and Herzog, 2010; Joumaa et al., 2008; Johnston et al., 2016; Leonard et al., 2010) to the whole-human level (Seiberl et al., 2015; Chapman et al., 2018; Chen et al. 2019), and is fundamental to a complete understanding of muscle contraction. Residual force enhancement (rFE) and residual force depression (rFD) are two history dependent force phenomena, characterized by increases and decreases in isometric force following active lengthening and shortening, respectively, in comparison to a reference isometric contraction at the same muscle length and level of activation (Abbott and Aubert, 1952; Chapman et al., 2018; Herzog, 2004; Rassier and Herzog, 2004b; Seiberl et al., 2015a). The magnitude of rFE is shown to be dependent on the amplitude of muscle lengthening and largely independent of lengthening velocity (Abbott and Aubert, 1952; Edman et al., 1978; Herzog and Leonard, 2002, 2005; Julian and Morgan, 1979; Fukutani et al., 2019b), while rFD appears to be strongly and positively related to the work (i.e., product of force × length change) of muscle shortening (Herzog and Leonard, 1997; Herzog et al., 2000; Lee et al., 2000). Suggested mechanisms include length-dependent contributions from non-contractile elements (i.e., titin) during lengthening for rFE (Herzog, 1998; Noble, 1992; Labeit et al., 2003; Nishikawa et al., 2012,; Joumaa and Herzog, 2014; Rode et al., 2009), and stress-induced angular actin deformations that impair cross bridge attachments during shortening for rFD (Joumaa et al., 2018; Maréchal and Plaghki, 1979). Given that rFE and rFD are likely to be intertwined in everyday movements (e.g., during stretch-shortening cycles), a means of maximizing rFE and/or minimizing rFD – the former being associated with increased neuromuscular economy (i.e., more force per unit of activation (Jones et al., 2016; Paquin and Power, 2018) and reduced adenosine triphosphate usage (Joumaa and Herzog, 2013) – would be desirable to improving motor efficiency.

While there have been recent investigations into the plasticity of the history dependence of force in states of altered contractile capacity (Dargeviciute et al., 2013; Power et al., 2012a, 2012b, 2013, 2014a, 2014b; Ramsey et al., 2010; de Ruiter et al., 2000; de Ruiter and de Haan, 2003), there is still little work looking into its trainability (Chen and Power, 2019; Siebert et al., 2016). Prior studies have demonstrated contraction-type-dependent morphological adaptations with training, whereby eccentric loading leads to increases in fascicle lengths/serial sarcomere numbers (Butterfield and Herzog, 2005; Franchi et al., 2014; Lynn and Morgan, 1994; Lynn et al., 1998; Reeves et al., 2009; Timmins et al., 2016) and concentric loading leads to decreases in fascicle lengths/serial sarcomere numbers (Butterfield and Herzog, 2005; Lynn and Morgan, 1994; Lynn et al., 1998, Timmins et al., 2016; Morais et al. 2019). Considering that rFE and rFD are length-dependent properties, eccentric- and concentric-biased training could potentially modify rFE and rFD through respective increases and decreases to the number of sarcomeres in series that modulate the amplitude of sarcomere lengthening and shortening experienced for a given muscle length change. A study by Chen and Power (2019) found that rFE (but not rFD) was differentially increased by chronic (4-week) concentric resistance training, and decreased by chronic eccentric resistance, owing to a likely combination of mechanical and neurological factors. While there was ultrasound evidence of fascicle length adaptations, the data were obtained from a small subset of the participants. As such, it remains to be elucidated: (i) what mechanical factors were responsible for the alterations to history-dependent force properties, and (ii) whether these alterations would persist after accounting for supposed serial sarcomere number differences.

Therefore, the purpose of these experiments was to gain additional mechanistic insight into the adaptability of the history dependence of force. We employed a model of uphill- and downhill-running rats to allow for an isolated (*in vitro*) approach to investigate muscle contractile properties and sarcomerogenesis. It was hypothesized that for an absolute muscle length change, rFE and rFD would increase following concentric training and decrease following eccentric training, owing to a respective reduction and addition of sarcomeres in series. However, should differences in rFE/rFD persist after normalizing for presumed serial sarcomere number discrepancies, mechanisms other than lengthening/shortening amplitudes must contribute to the modifiability of the history dependence of force following chronic incline/decline running.

## Methods

### Animals

Thirty-one male CD^®^ Sprague-Dawley IGS rats (sacrificial age: 18.8 ± 0.28 weeks, mass: 523.6 ± 11.0 g) were obtained (Charles River Laboratories, Senneville, QC, Canada) for study, with approval from the Animal Care Committee of the University of Guelph. Rats were housed in groups of three, with a maximum of two groups at any given time, and free fed a Teklad global 18% protein rodent diet (Envigo, Huntingdon, Cambs., UK) and water. After a week of acclimation to the new housing conditions, each rat was familiarized with running and assigned to one of three experimental groups: uphill running, downhill running, and sedentary control (i.e., no running intervention). Following the 20 days of exercise, rats recovered for 72 hours before sacrifice via CO_2_ asphyxiation followed by cervical dislocation, for experimental testing.

### Training protocol

One week prior to training, rats were familiarized with treadmill running (on a 0° grade). Rats in the exercise intervention groups (i.e., uphill/downhill running) were run 5 days/wk (i.e., Mon-Fri) on an EXER 3/6 animal treadmill (Columbus Instruments, Columbus, OH, USA) set to a 15° incline or decline for 20 training days, over a 4-week period. Training sessions lasted 15 minutes on the first day and the daily duration was increased by 5 min/day, up to the target 35 minutes (by the fifth training day) for the remainder of the training period. At the start of each training session, rats were first introduced to a walking speed of 10 m/min, which was gradually increased to the 16 m/min target running speed, at a rate of 1 m every min. Rats were provided with 2 minutes of rest after each 5 minute bout. Running was encouraged with a firm tap or push from behind. Introduction into the training protocol was staggered so that the subsequent group could begin training once the previous group was halfway through their training period.

### Experimental set up

Following sacrifice, the soleus (SOL) and extensor digitorum longus (EDL) muscles were carefully harvested from the right hind limbs. Silk-braided sutures (USP 2-0, metric 3) were tied along the musculotendinous junctions and mounted to the force-length controller/transducers in the 806D Rat Apparatus (Aurora Scientific, Aurora, ON, Canada). The muscles were bathed in a ~25°C Tyrode solution (121 mM NaCl, 24 mM NaHCO_3_, 5.5 mM D-Glucose, 5 mM KCl, 1.8 mM CaCl_2_, 0.5 mM MgCl_2_, 0.4 mM NaH_2_PO_4_, 0.1 mM EDTA) and bubbled with a 95% O_2_/5% CO_2_ gas mixture (Praxair Canada Inc., Kitchener, ON, Canada) to a pH of ~7.4. A 701C High-Powered, Bi-Phase Stimulator (Aurora Scientific, Aurora, ON, Canada) was used to evoke all contractions via two parallel platinum electrodes, submerged in the solution, situated on either side of the muscle. Force, length, and stimulus trigger data were all sampled at 10,000 Hz with a 605A Dynamic Muscle Data Acquisition and Analysis System (Aurora Scientific, Aurora, ON, Canada). All data were analyzed with the 615A Dynamic Muscle Control and Analysis High Throughput (DMC/DMA-HT) software suite (Aurora Scientific, Aurora, ON, Canada).

### Experimental procedures

An experimental schematic is depicted in Fig. 1. Testing with each muscle proceeded in order from Protocol A to G, with the order of fixed and relative protocols being randomized within the rFD and rFE conditions. All rFD and rFE protocols consisted of an rFD/rFE trial and a reference isometric (ISO) trial. The rFD trials (in protocol C and D) always preceded the ISO trials so as that the comparatively lower forces in the rFD trials could be attributed to shortening-induced deficits, as opposed to fatigue. Likewise, the ISO trials (in protocol E and F) always preceded the rFE trials so that the comparatively higher forces in the rFE trials could be attributed to lengthening-induced enhancements, rather than fatigue. Contractions in Protocols C to G were separated by 5 min intervals to minimize muscle fatigue. Prior to the start of the protocols, the muscle was passively set to a taut length (measured tie-to-tie) that exerted ~0.075 N of resting tension, as a starting point for approximating optimal lengths for tetanic force production (L_o_).

**Figure 1:**
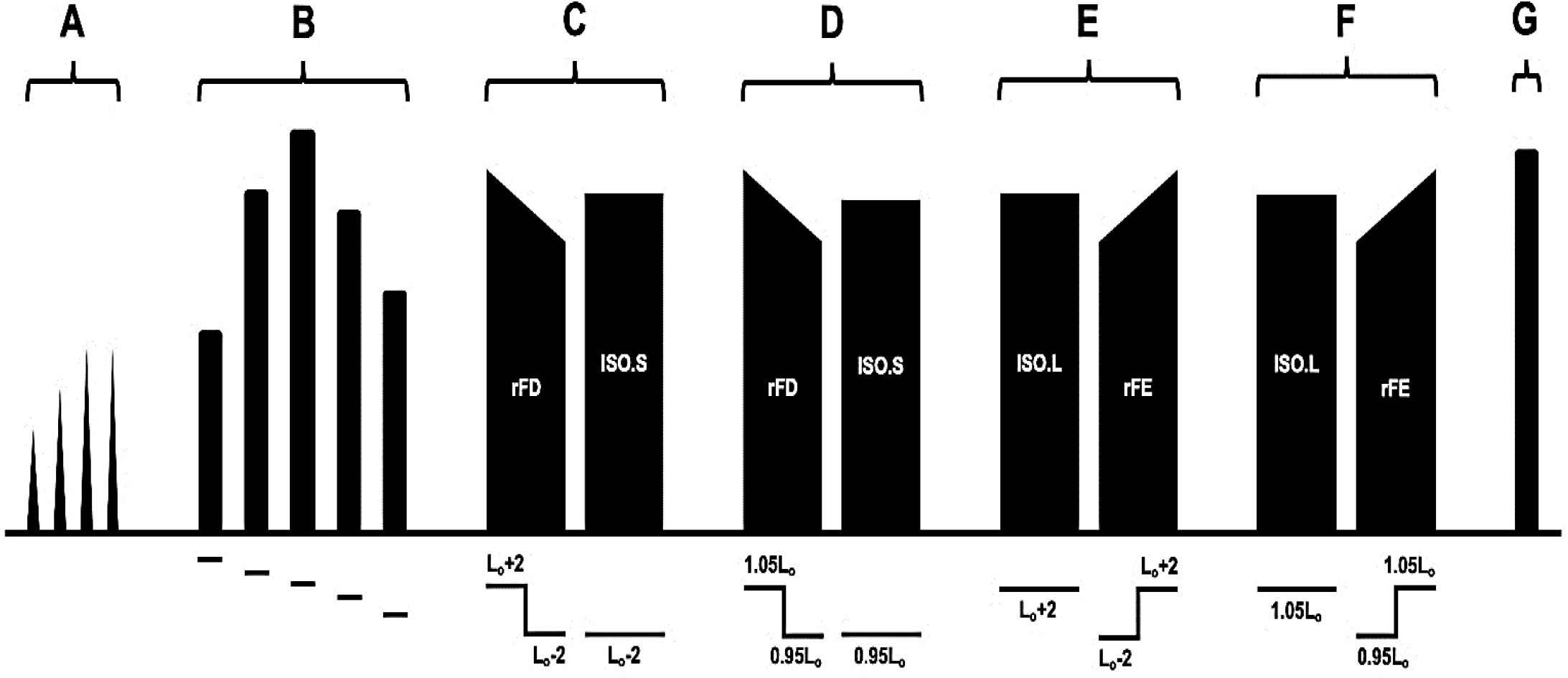
Schematic experimental timeline. **A.** Determination of twitch current. **B.** Determination of L_o_/F_o_. **C.** Fixed rFD protocol. **D.** Relative rFD protocol. **E.** Fixed rFE protocol. **F.** Relative rFE protocol. **G.** Final muscle assessment. The fixed and relative protocols were randomized. *L_o_, optimal muscle length; F_o_, isometric force at optimal muscle length; rFD, residual force depression; ISO.S, short reference isometric; ISO.L, long reference isometric; rFE, residual force enhancement*.

#### Protocol A: Twitch current

Single 1.25-ms pulses were incremented by 0.5 mA (starting from 1 mA) until a current suitable to elicit peak twitch force was determined.

#### Protocol B: Optimal length

A maximal tetanic stimulation (pulse width: 0.3 ms, duration: 1 s, frequency: 100 Hz) was delivered before the muscle was passively shortened or stretched by 2 mm and a subsequent stimulation was delivered 1.5 min later. This was repeated until peak tetanic force was obtained and the corresponding muscle length was measured. Once L_o_ was roughly determined, 1-mm and 0.5-mm increments were further used to more accurately establish L_o_.

#### Protocol C: Fixed rFD

The fixed rFD trial was comprised of 3 distinct yet continuous phases: a 1-s pre-activation at L_o_+2 mm, a −2 mm/s isokinetic shortening to L_o_-2 mm, and a 3-s isometric at L_o_-2 mm. The subsequent ISO.S trial consisted of a 6-s isometric at L_o_-2 mm.

#### Protocol D: Relative rFD

The relative rFD trial was comprised of 3 distinct yet continuous phases: a 1-s pre-activation at 1.05L_o_, a −2 mm/s isokinetic shortening to 0.95L_o_, and a 3-s isometric at 0.95L_o_. The subsequent ISO.S trial consisted of a 6-s isometric at 0.95L_o_.

#### Protocol E: Fixed rFE

The fixed rFE trial was comprised of 3 distinct yet continuous phases: a 1-s pre-activation at L_o_-2 mm, a +2 mm/s isokinetic lengthening to L_o_+2 mm, and a 3-s isometric at L_o_+2 mm. The preceding ISO.L trial consisted of a 6-s isometric at L_o_+2 mm.

#### Protocol F: Relative rFE

The relative rFE trial was comprised of 3 distinct yet continuous phases: a 1-s pre-activation at 0.95L_o_, a +2 mm/s isokinetic lengthening to 1.05L_o_, and a 3-s isometric at 1.05L_o_. The preceding ISO.L trial consisted of a 6-s isometric at 1.05L_o_.

#### Protocol G: Final muscle assessment

A final tetanic contraction was performed at L_o_ to assess decreases in isometric force-generating capacity.

#### Serial sarcomere number estimations

FolLowing mechanical testing, muscles were removed from the bath, passively stretched to L_o_ (determined from protocol B), tied to wooden sticks, and fixed in 10% phosphate-buffered formalin for 48 hrs, rinsed with phosphate-buffered saline, and digested in 30% nitric acid for at least 6 hrs to remove connective tissue and allow individual muscles fascicles to be teased out (Butterfield et al., 2005). Six fascicles were carefully obtained from deep and superficial regions of each muscle. Fascicle lengths (FL) were measured using 150 mm (resolution: 0.01 mm) electronic digital calipers (Marathon, Vaughan, ON, Canada). Sarcomere length (SL) measurements were taken over six different regions across the length of each fascicle via laser diffraction (Coherent, Santa Clara, CA, USA) with a 5 mW diode laser (beam diameter: 25 μm, wavelength: 635 nm) and custom LabVIEW program (Version 2011, National Instruments, Austin, TX, USA) (Lieber et al., 1984), for a total of 36 SL measurements per muscle. Serial sarcomere numbers (SSN) were calculated as:

> ***Serial sarcomere number = fascicle length / average sarcomere length***

### Data analyses

Steady-state isometric force values were calculated by subtracting passive force (resting baseline, pre-activation) from active force 2-s and 1-s following the end of the active length change for the SOL and EDL, respectively. Timing of ‘steady-state’ values were based on previous analysis of the rate of tension decay after stretch (Ramsey et al., 2010), in conjunction with visual confirmation from our own observations on the dissipation of force transients. Absolute rFE and rFD values (expressed in N/cm^2^) were calculated as the increase and decrease in isometric force following active lengthening and shortening, respectively, relative to the ISO contraction at the same time point into the active contraction (i.e., roughly 5-s and 4-s into the 6-s contraction for the SOL and EDL, respectively). Whereas, normalized rFE and rFD values (expressed as %ISO) were calculated as the percent difference in steady-state isometric force, relative to the ISO contraction, at the same time point. Similarly, passive FE (pFE) was calculated as the absolute and percent increase in passive force following active lengthening, relative to passive force (at rest, post-activation) of the ISO contraction. Peak eccentric force values were taken as the highest value achieved during the active lengthening phase of the rFE trials. Work of shortening values were calculated as the force-displacement integral for the active shortening phase of the rFD trials. All force and work values were normalized to physiological cross-sectional area (PCSA in cm^2^). PCSA was calculated as:

1. ***PCSA = muscle mass / (muscle density × normalized fascicle length)*** Wherein, *muscle density* was assumed to be 1.112 g/cm^3^ (Ward and Lieber, 2005), and normalized fascicle length was calculated as:
2. ***Normalizedfascicle length = fascicle length × (optimal sarcomere length / measured sarcomere length*)** Wherein, *optimal sarcomere length* was assumed to be 2.72 μm (based the average of all SL measurements)

Normalized fascicle length was used to calculate PCSA in order to account for inconsistencies in L_o_ that arose during the muscle fixation/digestion process.

A two-tailed paired *t*-test was used to compare force values between the history dependent and ISO trials to ascertain the presence of rFE, rFD, and pFE. A two-way analysis of variance (ANOVA) with a Holm-Sidak analysis for all pairwise comparisons was used to compare SSN, SL, FL, L_o_, and F_o_ values across muscles (SOL, EDL) and training groups (control, uphill, downhill). A three-way ANOVA with a Holm-Sidak analysis for all pairwise comparisons was used to compare peak eccentric force, work of shortening, rFD, rFE, and pFE values across muscles (SOL, EDL), training groups (control, uphill, downhill), and protocols (fixed, relative). Significance was set to an *α* = 0.05. All data are reported as means ± standard errors (SE).

## Results

### Muscle force and work

There was no effect of muscle (*F*(1,56) = 1.661, *p* = 0.203) or training (*F*(1,56) = 0.100, *p* = 0.905) for isometric force at optimal length (Table 1). There was a main effect of muscle (*F*(1,108) = 62.378, *p* < 0.001) for peak eccentric force, but no effect of training (*F*(1,108) = 0.126, *p* = 0.882) or protocol (*F*(1,108) = 0.895, *p* = 0.346), whereby the SOL produced 61% higher eccentric force than the EDL (Table 1). Furthermore, there was a main effect of muscle (*F*(1,108) = 19.043, *p* < 0.001) and protocol (*F*(1,108) = 4.167, *p* = 0.044) for work of shortening, but no muscle × protocol interaction (*F*(1,108) = 1.988, *p* = 0.161) or effect of training (*F*(1,108) = 0.0522, *p* = 0.949), such that work performed during shortening was 37% greater for the SOL than EDL and 15% greater for the fixed compared to relative protocol (Table 1).

**Table 1.** Force and work values across the various training groups and protocols for the SOL and EDL (*n* = 31 male rats). Isometric force was not significantly different across muscles (*p* = 0.203) or training groups (*p* = 0.905). Peak eccentric force was 61% greater for the SOL than EDL. Work of shortening was 37% greater for the SOL than EDL (*p* < 0.001) and 15% greater for the fixed compared to relative protocol (*p* = 0.044). Normalized values of isometric force, eccentric force, and work of shortening were 34%, 96%, and 65% greater for the SOL than EDL, respectively (*p* < 0.05).

|  | Soleus (SOL) |  |  | Extensor Digitorum Longus (EDL) |  |  |
| --- | --- | --- | --- | --- | --- | --- |
|  | Control | Uphill | Downhill | Control | Uphill | Downhill |
| Iso force, N | $1.39 \pm 0.07$ | $1.21 \pm 0.15$ | $1.23 \pm 0.17$ | $1.10 \pm 0.08$ | $1.21 \pm 0.08$ | $1.15 \pm 0.13$ |
| Iso force, N/cm<sup>2</sup>* | $9.51 \pm 0.87$ | $7.95 \pm 1.47$ | $9.03 \pm 1.31$ | $6.41 \pm 0.66$ | $6.95 \pm 0.87$ | $6.43 \pm 0.89$ |
| **Fixed Protocol** | Control | Uphill | Downhill | Control | Uphill | Downhill |
| Ecc force, N* | $2.44 \pm 0.12$ | $2.25 \pm 0.25$ | $2.29 \pm 0.19$ | $1.22 \pm 0.14$ | $1.47 \pm 0.12$ | $1.51 \pm 0.20$ |
| Ecc force, N/cm<sup>2</sup>* | $16.78 \pm 1.55$ | $14.38 \pm 2.23$ | $16.90 \pm 1.73$ | $7.01 \pm 0.91$ | $8.32 \pm 0.99$ | $8.46 \pm 1.31$ |
| Con work, mJ*† | $3.34 \pm 0.22$ | $2.99 \pm 0.41$ | $3.00 \pm 0.43$ | $1.96 \pm 0.18$ | $2.19 \pm 0.19$ | $2.23 \pm 0.28$ |
| Con work, mJ/cm<sup>2</sup>* | $23.10 \pm 2.46$ | $19.82 \pm 3.89$ | $22.06 \pm 3.48$ | $11.37 \pm 1.31$ | $12.68 \pm 1.90$ | $12.59 \pm 1.88$ |
| **Relative Protocol** | Control | Uphill | Downhill | Control | Uphill | Downhill |
| Ecc force, N* | $2.23 \pm 0.14$ | $2.12 \pm 0.29$ | $2.07 \pm 0.21$ | $1.26 \pm 0.16$ | $1.39 \pm 0.12$ | $1.52 \pm 0.22$ |
| Ecc force, N/cm<sup>2</sup>* | $15.48 \pm 1.71$ | $13.82 \pm 2.62$ | $15.37 \pm 1.91$ | $7.32 \pm 1.12$ | $7.86 \pm 1.04$ | $8.54 \pm 1.46$ |
| Con work, mJ*† | $2.76 \pm 0.19$ | $2.35 \pm 0.39$ | $2.47 \pm 3.57$ | $1.85 \pm 0.17$ | $2.12 \pm 0.27$ | $2.09 \pm 0.32$ |
| Con work, mJ/cm<sup>2</sup>* | $19.07 \pm 2.00$ | $15.69 \pm 3.63$ | $18.19 \pm 2.88$ | $10.77 \pm 1.29$ | $12.34 \pm 2.20$ | $11.92 \pm 2.04$ |

Values are means ± SE. \* Indicates an effect of muscle ( $p < 0.05$ ). † Indicates an effect of protocol ( $p < 0.05$ ). *Iso force*, isometric force at optimal muscle length; *Ecc force*, peak eccentric force; *Con work*, work of shortening.

In contrast, when normalized to PCSA, there was a main effect of muscle for specific (i.e., expressed in units/cm^2^) isometric force at optimal length (*F*(1,56) = 6.941, *p* = 0.011), specific peak eccentric force (*F*(1,108) = 65.911, *p* < 0.001), and specific work of shortening (*F*(1,108) = 27.885, *p* < 0.001), but no effect of training (*p* ≥ 0.05) or protocol (*p* ≥ 0.05). Specifically, the SOL exhibited 34% higher specific isometric force, 96% higher specific eccentric force, and performed 65% more specific work during shortening than the EDL across training groups and protocols (Table 1).

### Serial sarcomere numbers and muscle length measures

There was an interaction of muscle × training (*F*(2,371) = 3.237, *p* = 0.040) for SSN, in addition to main effects of muscle (*F*(1,371) = 119.567, *p* < 0.001) and training (*F*(2,371) = 5.732, *p* = 0.004). There was significantly greater SSN for the SOL than EDL (*p* = 0.001) across training groups and protocols, and the downhill group had a significantly greater SSN than both the uphill (*p* = 0.005) and control (*p* = 0.017) groups across muscles and protocols (Fig. 2). Within SOL, downhill running (95% CI = 6992.2, 7237.1) resulted in a 6% greater SSN (*p =* 0.003) than uphill running (95% CI = 6515.0, 6895.0), while SSN for the EDL did not significantly differ across training groups (*p* ≥ 0.05) (Fig. 2). Moreover, there was a main effect of muscle for fascicle length (*F*(1,56) = 39.358, *p* < 0.001) and muscle length (*F*(1,52) = 604.518, *p* < 0.001), but no effect of training (*p* ≥ 0.05) for either dependent variable, such that the SOL had a 17% shorter muscle but 13% longer fascicles compared to the EDL (Table 2). In contrast, there was no effect of either muscle (*F*(1,55) = 0.734, *p* = 0.395) or training (*F*(2,55) = 0.182, *p* = 0.834) on sarcomere lengths (Table 2).

**Figure 2:**
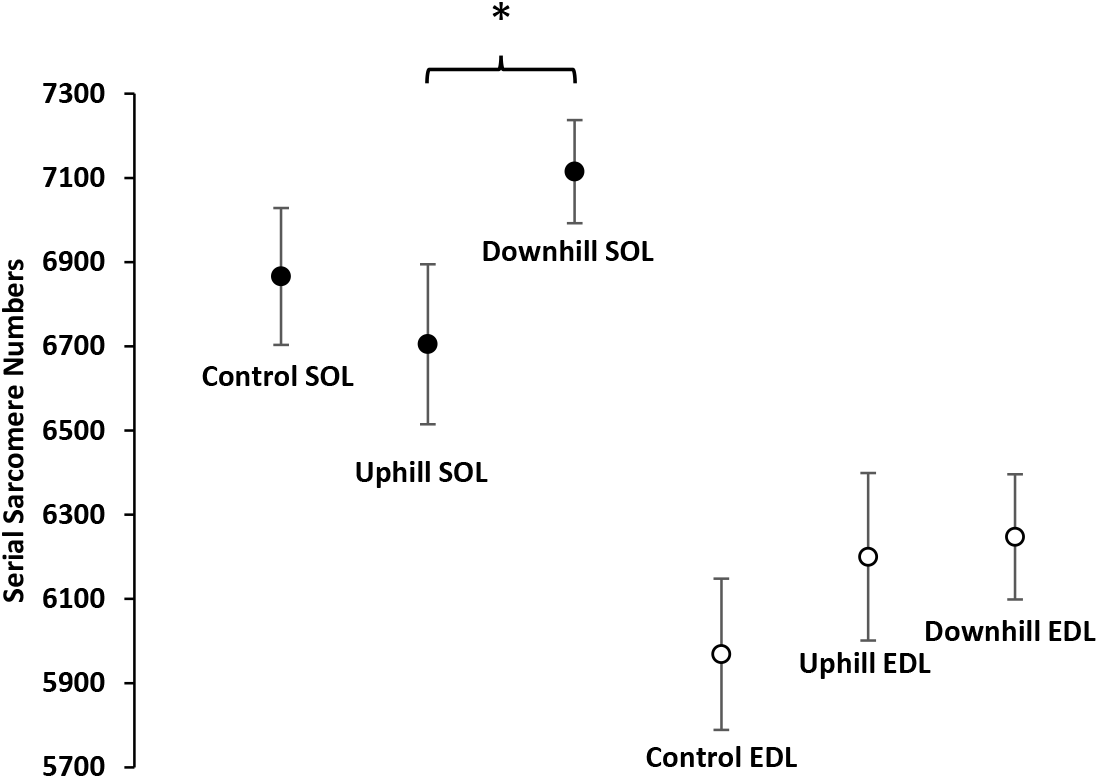
Sarcomerogenesis following uphill and downhill running. Mean ± 95% confidence intervals for SSN (*n* = 31 male rats). There was an interaction of muscle × training (*p* < 0.001) for SSN, in addition to main effects of muscle (*p* < 0.001) and training (*p* = 0.004). SSN within SOL were 6% greater for the downhill compared to uphill group (*p* = 0.003), and within the EDL there was a trend (*p* = 0.056) towards 4% more SSN for the downhill compared to control. * Indicates a significant difference (*p* < 0.05). *SSN, serial sarcomere numbers; SOL, soleus; EDL, extensor digitorum longus*.

**Table 2.** Muscle length measures across the various training groups for the SOL and EDL (*n* = 30 male rats). The SOL had a 17% shorter muscle (*p* < 0.001) but 13% longer fascicles (*p* < 0.001) compared to the EDL. However, there was no effect of training for either muscle length or fascicle length (*p* ≥ 0.05).

|  | Soleus (SOL) |  |  | Extensor Digitorum Longus (EDL) |  |  |
| --- | --- | --- | --- | --- | --- | --- |
|  | Control | Uphill | Downhill | Control | Uphill | Downhill |
| SL, $\mu\text{m}$ | $2.75 \pm 0.11$ | $2.76 \pm 0.14$ | $2.77 \pm 0.09$ | $2.75 \pm 0.05$ | $2.62 \pm 0.11$ | $2.69 \pm 0.11$ |
| FL, mm* | $18.60 \pm 0.32$ | $18.53 \pm 0.51$ | $19.33 \pm 0.26$ | $16.27 \pm 0.51$ | $16.89 \pm 0.54$ | $16.94 \pm 0.26$ |
| $L_o$, mm* | $30.75 \pm 0.86$ | $30.62 \pm 0.73$ | $31.08 \pm 0.51$ | $36.88 \pm 0.67$ | $37.27 \pm 0.86$ | $37.70 \pm 0.63$ |

Values are means ± SE. \* Indicates an effect of muscle ( $p < 0.05$ ). *SL*, average sarcomere length; *FL*, fascicle length; *Lo*, optimal muscle length.

### Residual ~force depression (Fig. 3A)

There was a main effect of muscle (*F*(1,108) = 11.892, *p* < 0.001) for absolute rFD (expressed in N/cm^2^), but no effect of training (*F*(2,108) = 0.440, *p* = 0.645) or protocol (*F*(1,108) = 2.793, *p* = 0.098). Absolute rFD was present across all training groups and protocols (*p* < 0.05) for both muscles, with 45% greater absolute rFD for the SOL than EDL (Fig. 3B). Absolute rFD was positively and linearly associated with specific work of shortening (Fig. 3D) for the SOL (R^2^ = 0.71; *F*(1,29) = 70.292, *p* < 0.001) and EDL (R^2^ = 0.61; *F*(1,29) = 45.362, *p* < 0.001). In contrast, there was no significant relationship between absolute rFD and SSN (SOL: R^2^ = 0.051, *p* = 0.228; EDL: R^2^ = 0.107, *p* = 0.073), fascicle length (SOL: R^2^ = 0.012, *p* = 0.565; EDL: R^2^ = 0.104, *p* = 0.077), or muscle length (SOL: R^2^ = 0.0004, *p* = 0.914; EDL: R^2^ = 0.113, *p* = 0.074) for either muscle.

**Figure 3:**
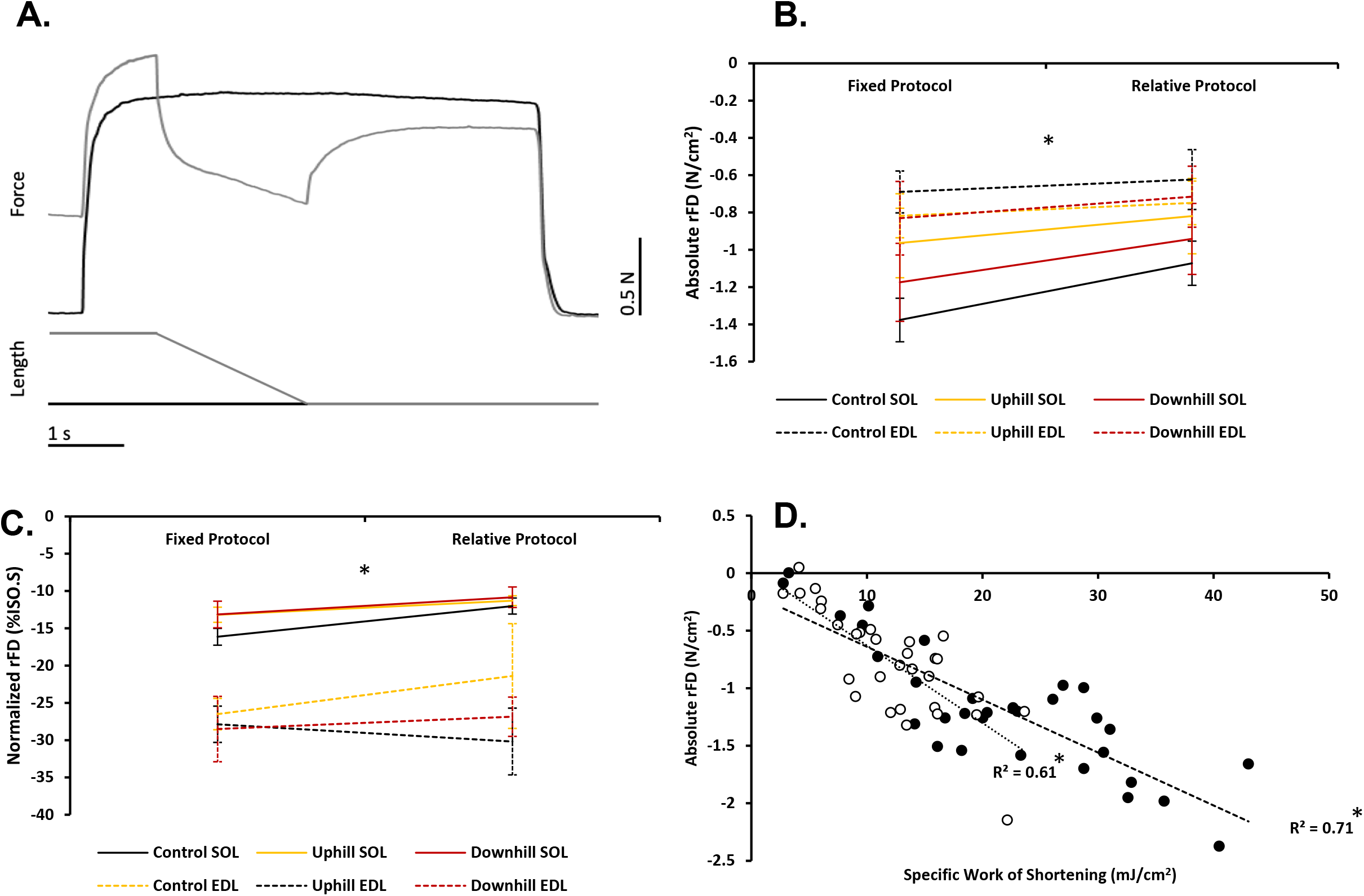
Residual force depression. **A.** Experimental data traces from the soleus, with the rFD contraction in gray and ISO.S contraction in black. Note the comparatively lower isometric force in the rFD condition for the same muscle length (and level of activation). **B.** Absolute rFD values across the various training groups and protocols for the SOL and EDL (*n* = 31 male rats). Absolute rFD was 45% greater for the SOL than EDL (*p* < 0.001) but was not significantly different across training groups (*p* = 0.645) or protocols (*p* = 0.098). **C.** Normalized rFD values across the various training groups and protocols for the SOL and EDL (*n* = 30 male rats). Normalized rFD was 2.1× greater (*p* < 0.001) for the EDL than SOL but was not significantly different across training groups (*p* = 0.745) or protocols (*p* = 0.219) (*n* = 31 male rats). **D.** Relationship between specific work of shortening and absolute rFD values across training groups during the fixed protocol for the SOL (black) and EDL (white) (*n* = 31 male rats). Specific work of shortening was positively and linearly associated with absolute rFD for both the SOL (R^2^ = 0.71, *p* < 0.001) and EDL (R^2^ = 0.61, *p* < 0.001). * Indicates a significant relationship (*p* < 0.05). Values are means ± SE. * Indicates an effect of muscle (*p* < 0.05). *ISO.S, short reference isometric; rFD, residual force depression; SOL, soleus; EDL, extensor digitorum longus*.

Likewise, there was a main effect of muscle (*F*(1,108) = 67.200, *p* < 0.001) for normalized rFD (expressed as %ISO.S), but no effect of training (*F*(2,108) = 0.295, *p* = 0.745) or protocol (*F*(1,108) = 1.526, *p* = 0.219). Normalized rFD was present across all training groups and protocols (*p* < 0.05) for both muscles, with 2.1× greater normalized rFD for the EDL than SOL (Fig. 3C). There was no significant relationship between normalized rFD and specific work of shortening (SOL: R^2^ = 0.028, *p* = 0.364; EDL: R^2^ = 0.026, *p* = 0.386), SSN (SOL: R^2^ = 0.087, *p* = 0.113; EDL: R^2^ = 0.001, *p* = 0.845), fascicle length (SOL: R^2^ = 0.051, *p* = 0.223; EDL: R^2^ = 0.001, *p* = 0.845), or muscle length (SOL: R^2^ = 0.001, *p* = 0.851; EDL: R^2^ = 0.113, *p* = 0.074) for either muscle.

### Residual force enhancement (Fig 4A)

There was a main effect of muscle (*F*(1,08) = 30.138, *p* < 0.001) and protocol (*F*(1,2) = 0.0736, *p* = 0.004) for absolute rFE (expressed in N/cm^2^), but no significant muscle × protocol interaction (*F*(2,108) = 3.908, *p* = 0.051) or effect of training (*F*(2,108) = 0.0736, *p* = 0.929). Absolute rFE was present across all training groups and protocols (*p* < 0.05) for both muscles (Fig. 4B), wherein absolute rFE was 2.3x greater for the SOL than EDL across training groups and protocols, and 56% greater for the fixed compared to relative protocol across muscles and training groups. Absolute rFE was positively and linearly associated with specific eccentric force (Fig. 4D) for the SOL (R^2^ = 0.57; *F*(1,29) = 39.108, *p* < 0.001) and EDL (R^2^ = 0.72; *F*(1,29) = 76.100, *p* < 0.001). Whereas, there was no significant relationship between absolute rFE and SSN (SOL: R^2^ = 0.0004, *p* = 0.913; EDL: R^2^ = 0.086, *p* = 0.110), fascicle length (SOL: R^2^ = 0.004, *p* = 0.744; EDL: R^2^ = 0.079, *p* = 0.127), or muscle length (SOL: R^2^ = 0.012, *p* = 0.573; EDL: R^2^ = 0.001, *p* = 0.856) for either muscle.

**Figure 4:**
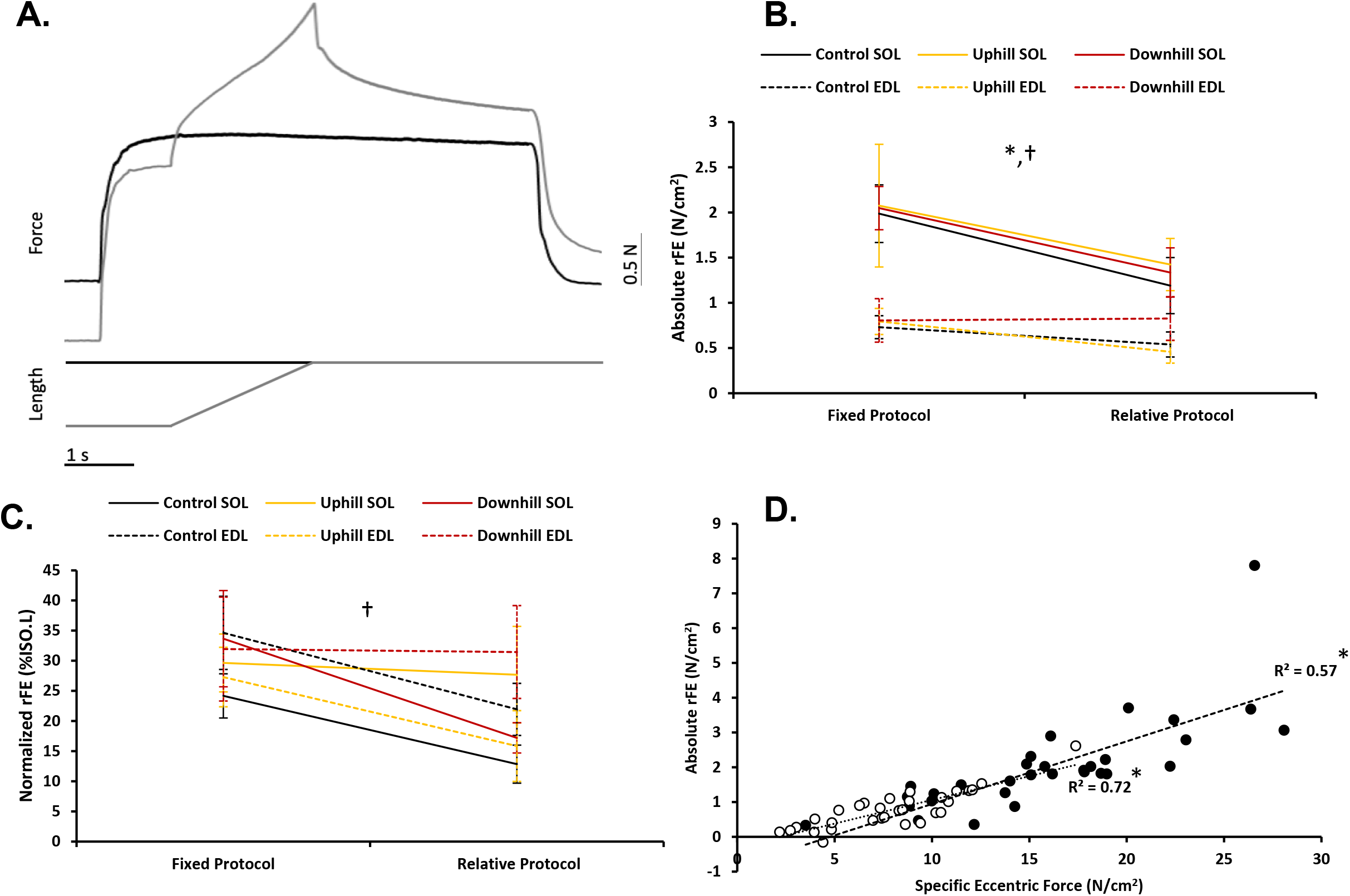
Residual force enhancement. **A.** Experimental data traces from the soleus, with the rFE contraction in gray and ISO.L contraction in black. Note the comparatively higher isometric force in the rFE condition (and comparatively higher passive force in the pFE condition) for the same muscle length (and level of activation). **B.** Absolute rFE values across the various training groups and protocols for the SOL and EDL (*n* = 31 male rats). Absolute rFE was 2.3× greater for the SOL than EDL (*p* < 0.001) and 56% greater for the fixed compared to relative protocol (*p* = 0.004), with no significant difference across training groups (*p* = 0.929). **C.** Normalized rFE values across the various training groups and protocols for the SOL and EDL (*n* = 30 male rats). Normalized rFE was 43% greater for the fixed compared to relative protocol (*p* = 0.009) but was not significantly different across muscles (*p* = 0.386) or training groups (*p* = 0.450). **D.** Relationship between specific eccentric force and absolute rFE values across training groups during the fixed protocol for the SOL (black) and EDL (white) (*n* = 31 male rats). Specific eccentric force was positively and linearly associated with absolute rFE for both the SOL (R^2^ = 0.57, *p* < 0.001) and EDL (R^2^ = 0.73, *p* < 0.001). *rFE, residual force enhancement; ISO.L, long reference isometric; pFE, passive force enhancement*. Values are means ± SE. * Indicates an effect of muscle (*p* < 0.05). † Indicates an effect of protocol (*p* < 0.05).

In contrast, there was a main effect of protocol (*F*(1,108) = 7.014, *p* = 0.009) for normalized rFE (expressed as %ISO.L), but no effect of muscle (*F*(1,108) = 0.759, *p* = 0.386) or training (*F*(2,108) = 0.805, *p* = 0.450). Normalized rFE was present across all training groups (*p* < 0.05) for both muscles, with 43% greater normalized rFE for the fixed compared to relative protocol (Fig. 4C). There was no significant relationship between normalized rFE and specific eccentric force (SOL: R^2^ = 0.026; *F*(1,29) = 0.852, *p* = 0.364; EDL: R^2^ = 0.022; *F*(1,29) = 0.641, *p* = 0.430), SSN (SOL: R^2^ = 0.054, *p* = 0.216; EDL: R^2^ = 0.039, *p* = 0.284), fascicle length (SOL: R^2^ = 0.040, *p* = 0.280; EDL: R^2^ = 0.040, *p* = 0.280), or muscle length (SOL: R^2^ = 0.004, *p* = 0.747; EDL: R^2^ = 0.006, *p* = 0.689) for either muscle.

### Passive force enhancement

There was a main effect of muscle (*F*(1,107) = 24.172, *p* < 0.001) for absolute pFE (expressed in N/cm^2^), but no effect of training (*F*(1,107) = 1.151, *p* = 0.320) or protocol (*F*(1,107) = 1.1491, *p* = 0.225). Absolute pFE was present across all training groups and protocols (*p* < 0.05) for both muscles, with 2.2× greater absolute pFE for the SOL than EDL (Fig. 5A). Absolute pFE was positively and linearly associated with absolute rFE (Fig. 5C) for the SOL (R^2^ = 0.41; *F*(1,29) = 19.897, *p* < 0.001) and EDL (R^2^ = 0.69; *F*(1,29) = 65.279, *p* < 0.001), amounting to 29% (0.71 ± 0.10 N/cm^2^) and 37% (0.30 ± 0.047 N/cm^2^) of the absolute rFE observed for the SOL and EDL, respectively. Furthermore, absolute pFE was positively and linearly associated with specific eccentric force (Fig. 5D) for the SOL (R^2^ = 0.49; *F*(1,29) = 28.212, *p* < 0.001) and EDL (R^2^ = 0.65; *F*(1,29) = 53.236, *p* < 0.001). Meanwhile, for the SOL, there was no significant relationship between absolute pFE and SSN (R^2^ = 0.010, *p* = 0.596), fascicle length (R^2^ = 0.032, *p* = 0.341), or muscle length (R^2^ = 0.124, *p* = 0.061). Whereas for the EDL, there was weak but significant relationship between absolute pFE vs. SSN (R^2^ = 0.158; *F*(1,29) = 5.458; *p* = 0.027) and absolute pFE vs. fascicle length (R^2^ = 0.150; F(1,29) = 5.124, *p* = 0.031), but not between absolute pFE and muscle length (R^2^ = 0.003, *p* = 0.768).

**Figure 5:**
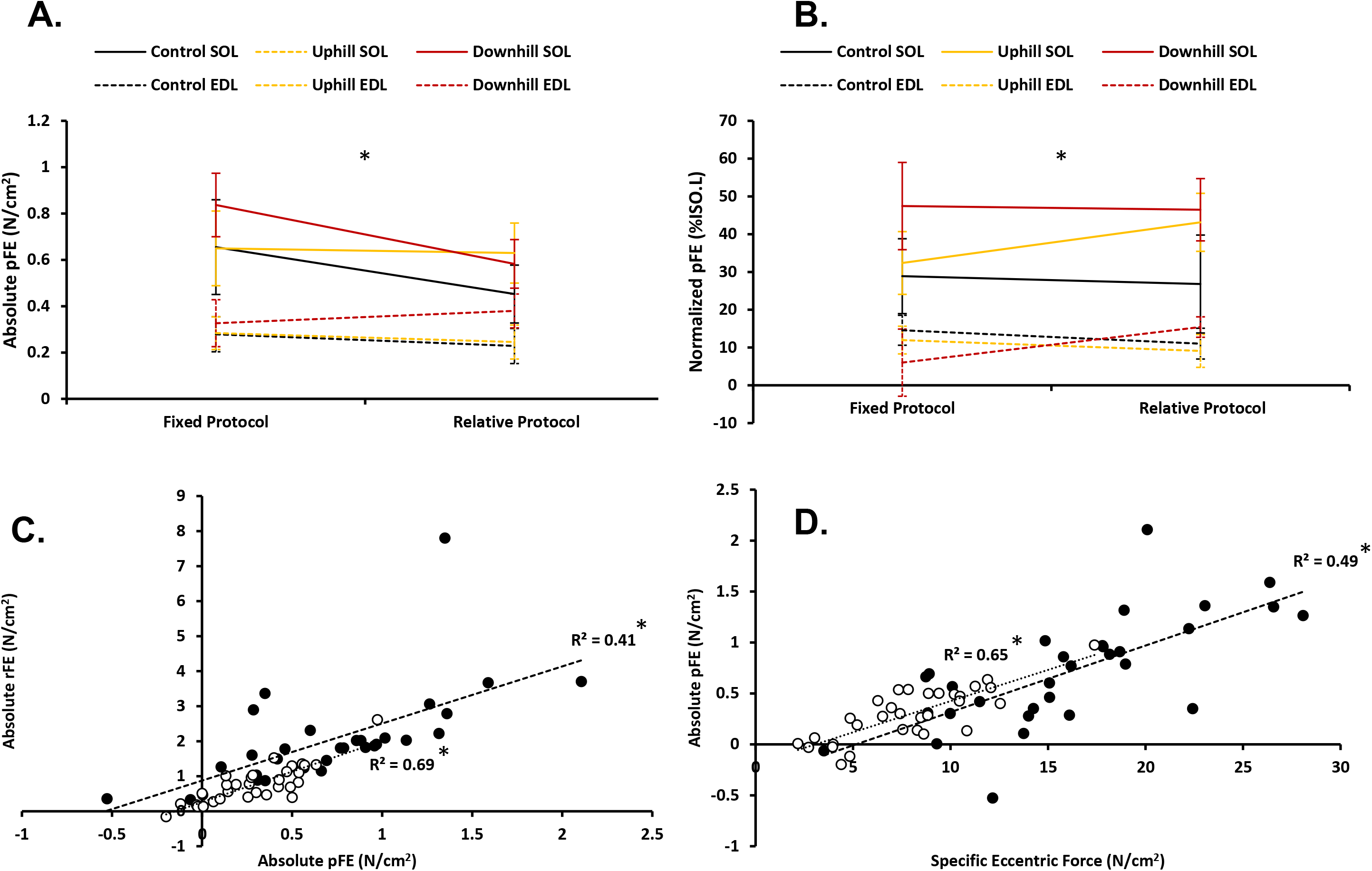
Passive force enhancement. **A.** Absolute pFE values across the various training groups and protocols for the SOL and EDL (*n* = 31 male rats). Absolute pFE was 2.2× greater for the SOL than EDL (*p* < 0.001) but was not significantly different across training groups (*p* = 0.320) or protocols (*p* = 0.225). **B.** Normalized pFE values across the various training groups and protocols for the SOL and EDL (*n* = 30 male rats). Normalized pFE was 3.3× greater for the SOL than EDL (*p* < 0.001) but was not significantly different across training groups (*p* = 0.320) or protocols (*p* = 0.700). **C.** Relationship between absolute pFE and absolute rFE values across training groups during the fixed protocol for the SOL (black) and EDL (white) (*n* = 31 male rats). Absolute pFE was positively and linearly associated with absolute rFE for both the SOL (R^2^ = 0.41, *p* < 0.001) and EDL (R^2^ = 0.69, *p* < 0.001). **D.** Relationship between specific eccentric force and absolute pFE values across training groups during the fixed protocol for the SOL (black) and EDL (white) (*n* = 31 male rats). Specific eccentric force was positively and linearly associated with absolute pFE for both the SOL (R^2^ = 0.49, *p* < 0.001) and EDL (R^2^ = 0.65, *p* < 0.001). Values are means ± SE. * Indicates an effect of muscle (*p* < 0.05). *pFE, passive force enhancement; SOL, soleus; EDL, extensor digitorum longus*.

Similarly, there was a main effect of muscle (*F*(1,107) = 31.979, *p* < 0.001) for normalized pFE (expressed as %ISO.L), but no effect of training (*F*(1,107) = 1.152, *p* = 0.320) or protocol (*F*(1,107) = 0.149, *p* = 0.700). Normalized pFE was present across all training groups and protocols (*p* < 0.05) for both muscles, with 3.3× greater normalized pFE for the SOL than EDL (Fig. 5B). There was no significant relationship between normalized pFE and normalized rFE for the SOL (R^2^ = 0.028, *p* = 0.367) but a significant, albeit weak, relationship for the EDL (R^2^ = 0.152; *F*(1,29) = 5.195, *p* = 0.030). Additionally, there was a weak but significant relationship between normalized pFE and specific eccentric force for the SOL (R^2^ = 0.180; *F*(1,29) = 6.377, *p* = 0.017) and EDL (R^2^ = 0.223; *F*(1,29) = 8.300, *p* = 0.007). Finally, there was no significant relationship between normalized pFE and SSN (SOL: R^2^ = 0.0003, *p* = 0.927; EDL: R^2^ = 0.050, *p* = 0.225), fascicle length (SOL: R^2^ = 3.644 × 10^−5^, *p* = 0.974; EDL: R^2^ = 0.047, *p* = 0.240), or muscle length (SOL: R^2^ = 0.036, *p* = 0.327; EDL: R^2^ = 0.009, *p* = 0.617) for either muscle.

## Discussion

The purpose of these experiments were to determine whether the history dependence of force could be modified by contraction-type-dependent SSN adaptations. Serial sarcomere number and history-dependent forces for the SOL and EDL muscles of rats were compared across training groups following 4-weeks of uphill and downhill running. In accordance with our hypothesis, there was a greater SSN for the SOL with downhill compared to uphill running, and a trend towards a greater SSN for the EDL with downhill running in comparison to the control group. While SSN appeared to differ with training, History-dependent forces were not similarly affected. Together, these results indicate that training-induced SSN adaptations do not modify whole-muscle *in-vitro* measures of rFD, rFE, and pFE in the lower hind limb muscles of rats.

### Serial sarcomere number adaptations

In the present study, SSN (Fig. 2) were consistent with the ~7500 count for the VL (Butterfield et al., 2005), with values across muscles ranging from ~3400 for the VI of rats (Lynn and Morgan, 1994; Lynn et al., 1998; Butterfield et al., 2005) to ~21,000 for the TA of rabbits (Butterfield and Herzog, 2006). While our measured 2.72 μm SL average was towards the higher end of the typical 2.4-2.8 μm range (Stephenson and Williams, 1982), values from this study were obtained from muscles (fascicles) passively fixed at their optimal length for force production, as opposed to given joint angle or skinned fibre. As a result, the compliance of the in series connective tissues would result in a shortening of sarcomeres upon activation and lengthening upon deactivation, and subsequently, an overestimation of SL when measured at a passive length.

In line with previous findings (Lynn and Morgan, 1994; Lynn et al., 1998; Butterfield et al., 2005; Morais et al. 2019), downhill running resulted in 6% greater SSN for the SOL when compared to uphill running, but not a sedentary control group. Moreover, while SSN were not significantly different across training groups for the EDL, there was a trend (*p* = 0.056) towards 4% greater SSN for the downhill vs. control group, indicating a potential stimulus for sarcomerogenesis with downhill running. Given that strain dynamics were not measured during training, insight into the actions of the SOL and EDL during uphill and downhill running have to be inferred based on previous observations. In a similar treadmill training program as the current study, Morais et al (2019) showed, in mice, concentric training led to a decrease in SSN for the VL and TA while eccentric training had no effect. Additionally, there was no training induced sarcomerogenesis effect for the GM and VM. However, when inferring SSN adaptations it is important to consider the specific muscle fibre excursion during locomotion (Hu et al. 2017). It is possible those muscles which did not show sarcomerogenesis did not experience a minimal threshold to warrant adaptations.

During rat locomotion, soleus activity begins just prior to foot contact and ends immediately before foot lift-off, whereas EDL activity begins just after foot lift-off and ends immediately after foot contact (Nicolopoulos-Stournaras and Iles, 1984). This would imply that the SOL is loaded eccentrically during downhill running and concentrically during uphill running, which would support the findings of increased SSN with eccentric training and decreased SSN with concentric training (Butterfield et al., 2005; Butterfield and Herzog, 2006). While not statistically significant, the greater apparent number of sarcomeres for both training groups in comparison to the sedentary control suggests that the EDL performed eccentric contractions during downhill and uphill running, which may be explained by eccentric braking actions at the onset of foot contact.

### Strength measures

Perhaps contrary to expectations, training did not alter isometric forces, eccentric forces, and work of shortening for either muscle. However, unlike the strength-oriented resistance training employed by Chen and Power (2019), the main goal of our incline/decline-running model was to induce differential SSN adaptations, which has previously been demonstrated for the quadriceps muscles of rats (Lynn and Morgan, 1994; Lynn et al., 1998; Butterfield et al., 2005) and is partially supported by the present findings for the lower hind limb muscles. Yet, SSN increases were not mirrored by increases in muscle length (Table 2) or force/work performed (Table 1). However, architectural adaptations at the level of the fascicle do not always translate to the whole muscle (Sharifnezhad et al., 2014), owing to factors such as pennation angles and connective tissue compliance, which mediate the transfer of forces across functional scales. Furthermore, there is the ever-present issue of ‘testing specificity’; assuming that neuromuscular adaptations are geared towards specific environmental perturbations, testing the muscle under different physiological/contractile conditions (i.e., *in vitro*) could likely wash out potential *in-vivo* responses (e.g., strain of specific muscle fibre regions during locomotion). Additionally, while SSN were significantly higher for the SOL following downhill vs. uphill running, from a functional standpoint, a 6% increase might not be sufficient to elicit prominent length-dependent changes in muscle force, as evidenced by the current results.

With respect to the rFE and rFD contractions, during a 4 mm muscle length change (i.e., in the fixed protocols) the 6% SSN difference for the SOL would roughly equate to an average sarcomere length change of 0.563 μm for the downhill group and 0.596 μm for the uphill group. However, this 0.033 μm difference in sarcomere length does not account for confounding variables such as pennation angle, curvature of the fascicle, connective tissue compliance, and sarcomere length non-uniformities. Therefore, irrespective of whether SSN adaptations also took place for the EDL, it is likely that SSN differences would not be large enough to promote length-dependent differences in force at the whole-muscle level. While it could be argued that longer training periods might lead to greater adaptations, Lynn and Morgan (1994) found that SSN counts peaked after just 1-week of incline/decline running, with no further changes leading up to the end of the 3-week training period. In a similar treadmill training program to the current study, Morais et al (2019) showed that, in mice, concentric training led to a decrease in SSN for the VL and TA while eccentric training had no effect. Additionally, there was no training induced sarcomerogenesis effect for the GM and VM. However, it remains to be determined how steeper inclines/declines, or specific active lengthening/shortening of a target muscle might affect training adaptations.

### Residual fforce depression

In line with the 8-72% rFD previously reported *in vitro* (Abbott and Aubert, 1952; Edman, 1975; Herzog and Leonard, 1997; Herzog et al., 1998; Joumaa and Herzog, 2010; Joumaa et al., 2012; Maréchal and Plaghki, 1979; Meijer, 2002; Morgan et al., 2000; Pun et al., 2010; Sugi and Tsuchiya, 1988), steady-state isometric force values following active shortening were 13% and 28% lower than the ISO for the SOL and EDL, respectively. Normalized rFD was 2.1× greater for the EDL (which is comprised of 95-100% type II fibres) than the SOL (which is comprised of 75-80% type I fibres) (Eng et al., 2008; Ranatunga and Thomas, 1990; Wigston and English, 1991), and is consistent with findings of 1.8× greater normalized rFD for type II compared to type I fibres from rabbit psoas and soleus muscle (Joumaa et al., 2015). On the other hand, absolute rFD was 45% greater for the SOL than EDL, contrary to the observations for normalized rFD. The discrepancy between relative and absolute values of rFD in the present study is most likely due to reduced contractile capacity (i.e., possibly fatigue) of the predominantly fast-twitch EDL muscle (Brooks and Faulkner, 1991), as evidenced by the comparatively lower overall strength measures (Table 1). Given that the underlying factor contributing to rFD is a stress-induced impairment of cross bridge attachments during active shortening, it is not surprising then that a reduced capacity to perform work during shortening would result in lower absolute values of rFD (Fig. 3D). However, after accounting for differences in isometric strength (i.e., lower contractile capacity), normalized rFD was greater for the EDL, suggesting greater intrinsic rFD for fast-twitch muscles, owing to distinct force-velocity relationships that facilitate the capacity to more produce work at a given shortening velocity, as compared with slow-twitch muscles (Joumaa et al., 2015; Pinnell et al. 2019).

Contrary to our hypothesis, absolute and normalized rFD values were not different following uphill and downhill training for either muscle, despite differences in SSN for the SOL, and possibly, EDL. As a result, it is possible that SSN adaptations were not prominent enough or specifically geared to evoke functional length-dependent differences under our *in-vitro* contractile conditions. The lack of training-induced alterations to rFD in the present study is consistent with findings from Chen and Power (2019), suggesting that neither SSN adaptations (in the present study) nor potential neurological adaptations (in Chen and Power, 2019) serve to modify rFD at the whole-muscle level.

### Residual force enhancement

In line with the 8-52% rFE previously reported *in vitro* (Abbott and Aubert, 1952; Edman et al., 1982; Meijer, 2002; Pun et al., 2010; Rassier and Pavlov, 2012; Sugi and Tsuchiya, 1988), steady-state isometric force values following active lengthening were 24% and 27% higher than the ISO for the SOL and EDL, respectively. Absolute rFE was 2.3× greater for the SOL than EDL while normalized rFE was not significantly different between muscles, contrary to observations from Ramsey et al. (2010) of ~55% greater normalized rFE for the EDL compared to SOL muscle of rats. In argument against reports that absolute rFE is preserved in conditions of reduced contractile force (Fukutani and Herzog, 2018b; Rassier and Herzog, 2004a), we show that absolute rFE is strongly and positively associated with the force established during the active lengthening phase of the rFE contraction (Fig. 4D), suggesting a strength-dependent component of rFE.

Although rFE is typically attributed to non-contractile elements (i.e., titin), contractile contributions (e.g., decreased cross bridge detachment) cannot be discounted (Rassier and Herzog, 2004a; Lee et al., 2007; Pinnell et al., 2019). Furthermore, cross bridge cycling is necessary for the engagement of both contractile and non-contractile elements involved in rFE and pFE (Fukutani et al., 2019b, Lee et al., 2007; Pinnell et al., 2019; Rassier and Herzog, 2005; Rassier et al., 2003; Herzog and Leonard, 2002; Rassier and Herzog, 2004a). Therefore, with reduced contractile capacity, cross bridge kinetics would become impaired, leading to hypothetical decreases in cross-bridge- and/or titin-mediated force enhancement. As such, the lower absolute rFE for the EDL (relative to the SOL) is attributed to a lower contractile capacity of muscle (Brooks and Faulkner, 1991), as evidenced by the comparatively lower overall strength measures for the EDL (Table 2). Conversely, after accounting for differences in isometric force (i.e., strength/weakness), normalized rFE was not different between the SOL and EDL, suggesting a lack of intrinsic difference for rFE between slow-twitch and fast-twitch muscles. While in contrast to Ramsey et al. (2010), and reports of stiffer titin isoforms for predominantly fast-twitch muscles (Horowits et al., 1986; Prado et al., 2005), observations from Fukutani et al. (2018b) of similar/greater absolute and normalized rFE values for the soleus (94.5% type I fibres) compared to psoas (100% type IIa fibres) muscle of rabbits (Prado et al., 2005) are in line with our findings.

Contrary to our hypothesis, absolute and normalized rFE values were not different following uphill and downhill training for either muscle, despite differences in SSN for the SOL, and possibly, EDL. In the same context as rFD, it is possible that the SSN adaptations were not prominent enough or specifically geared to evoke functional length-dependent differences under our *in-vitro* contractile conditions. The lack of training-induced alterations to rFE in the present study is in agreement with Siebert et al. (2015), but in contrast to findings from Chen and Power (2019), which reported increases to rFE with concentric training and decreases to rFE with eccentric training. Discrepancies between this study and the one by Chen and Power (2019) may be due to differences in training protocols, neurological components, and study designs. Training in Chen and Power (2019) was predominantly load-focused, with *in-vivo* torque measurements performed in the presence of an intact neurological system, the latter of which is primarily responsible for strength adaptations (e.g., increased motor unit activation/decreased neural inhibition) within the early phases (i.e., first 3-4 weeks) of resistance training (Douglas et al., 2017; Hahn, 2018). In contrast, the current training protocol involved a large aerobic component, with *in-vitro* force measurements performed independent of neurological contributions. Additionally, Chen and Power (2019) looked at within-individual comparisons (pre- and post-training), whereas, this study and the one by Siebert et al. (2016) compared values in a cross-sectional manner, which could have masked more minute responses. Ultimately though, the results suggest that length-dependent alterations to history-dependent properties via SSN adaptations do not serve to modify rFE (or rFD) at the whole-muscle level.

### Passive force enhancement

In line with the ~3-54% pFE previously observed in the literature (Herzog and Leonard, 2002; 2005; Lee and Herzog, 2002; Lee et al., 2007), resting forces following active lengthening were 37% and 11% higher than those from the ISO for the SOL and EDL, respectively. Additionally, pFE contributed to 29% and 37% of the rFE for the SOL and EDL, respectively, which is consistent with the ~8-84% pFE-to-rFE ratio reported in the literature (Herzog and Leonard, 2002; 2005; Lee and Herzog, 2002; Lee et al., 2007), and highlights a common history dependent mechanism. Considering that pFE is strongly and positively associated with rFE (Fig. 5C), the simultaneously greater absolute and normalized pFE values for the SOL compared to EDL is in contrast to the greater absolute but similar normalized rFE observed. Disparity between normalized values of rFE and pFE is most likely owing to differences between the isometric reference contractions, wherein active force (for rFE) is diminished but passive tension (for pFE) is preserved with lower contractile capacity. In other words, strength-related impairments to cross bridge cycling would lead to concurrent decreases in absolute rFE and pFE for the EDL (as illustrated by the strong, positive relationships to eccentric force in Fig. 4D and Fig. 5D), but normalization to similar resting forces would not amplify pFE values like smaller ISO forces would to rFE. As a consequence, comparisons cannot be made regarding intrinsic differences in pFE between the SOL and EDL from this study.

Although absolute values of pFE were not associated with SSN or fascicle lengths for the SOL, there were significant, albeit weak, relationships for the EDL, which highlight the length-dependence of pFE for fast-twitch muscles (shown to possess stiffer titin isoforms). Yet, the fact that absolute and normalized pFE values were not different following uphill and downhill training for either muscle in the present study only further reinforces the idea that SSN adaptations were ineffective at inducing *in-vitro*, length-dependent alterations to history-dependent forces at the whole-muscle level.

This is the first study to demonstrate SSN differences for the soleus, and possibly EDL, muscle of uphill- and downhill-running rats, which had previously only been reported for the VI, VL, and TA, and recently the VL: and TA of mice (Morais et al. 2019). Consequently, it appears that rFD, rFE, and pFE are not differentially modifiable at the whole-muscle level, on the basis of training-induced alterations to SSN, as a means of altering sarcomere shortening/lengthening amplitudes. However, these findings highlight the relevance of reporting both absolute and relative history-dependent force values, which suggest that absolute values of rFD, rFE, and pFE are not maintained during conditions of reduced contractile force, most likely owing to impaired cross bridge engagement. In addition to furthering our understanding of muscle contractions, from a practical standpoint, increases to rFE/decreases to rFD could serve to improve the neuromuscular economy of functional movements, particularly during SSCs, wherein rFE appears to attenuate rFD in an amplitude-dependent manner. As such, increases to rFE could benefit force production during both lengthening- and shortening-induced conditions. Considering how intrinsic cross bridge dynamics are to history dependent force properties (and vice versa), future studies investigating the trainability of the history dependence of force should look to improve the contractile capability of muscle through strength-focused training protocols.

## Acknowledgments

This project was supported by the Natural Sciences and Engineering Research Council of Canada (NSERC). Infrastructure was provided by the University of Guelph start-up funding. No conflicts of interest, financial or otherwise, are declared by the authors.

## Disclosure statement

No conflicts of interest, financial or otherwise, are declared by the authors.

## Ethics statement

All procedures were approved by the Animal Care Committee of the University of Guelph.

## Data accessibility

Individual values of all supporting data are available upon request.

## Grants

This project was supported by the Natural Sciences and Engineering Research Council of Canada (NSERC). Infrastructure was provided by the University of Guelph start-up funding.

## Author contributions

All authors contributed equally

